# Working memory as a representational template for reinforcement learning

**DOI:** 10.1101/2024.04.25.591119

**Authors:** Kengo Shibata, Verena Klar, Sean J Fallon, Masud Husain, Sanjay G Manohar

**Affiliations:** Nuffield Department of Clinical Neurosciences, University of Oxford, Oxford OX3 9DU, UK; Department of Experimental Psychology, University of Oxford, Oxford OX2 6GG, UK; School of Psychology, University of Plymouth, Plymouth PL4 8AA, UK

**Keywords:** Working memory, reinforcement learning, reversal learning, feature-based representations, object-based representations, representational templates

## Abstract

Working memory (WM) and reinforcement learning (RL) both influence decision-making, but how they interact to affect behaviour remains unclear. We assessed whether RL is influenced by the format of visual stimuli in WM, either feature-based or unified, object-based representations. In a pre-registered paradigm, participants learned stimulus-action combinations, mapping four stimuli onto two feature dimensions to one of two actions through probabilistic feedback. In parallel, participants retained the RL stimulus in WM and were asked to recall this stimulus after each trial. Crucially, the format of representation probed in WM was manipulated, with blocks encouraging either separate features or bound objects to be remembered. Incentivising a feature-based WM representation facilitated feature-based learning, shown by an improved choice strategy. This reveals a role of WM in providing sustained internal representations that are harnessed by RL, providing a framework by which these two cognitive processes cooperate.

## Introduction

Reinforcement learning (RL) is a key process through which biological and artificial agents incrementally learn actions by interacting with the environment and receiving iterative feedback^1^. Credit assignment to the relevant attribute is a key aspect for successful learning for humans and artificial agents alike. For instance, consider sending an email to a collaborator. Unexpectedly, you get a very quick reply, and you would like to reinforce the action that led to such a quick response. You might infer that it was the time you sent it, or the formatting, or the subject line. Which strategy you end up reinforcing might depend on what is currently on your mind. Humans flexibly *assign credit* to different features of the world, depending on what they pay attention to. This process has been behaviourally tested using the Wisconsin Card Sorting Test^2^, where subjects learn to sort multi-featured cards, inferring the relevant features based on reward feedback. A major open question for this process is how the brain determines the set of states on which reinforcement operates. Here, we propose and test a new framework in which working memory (WM) plays a central role in providing a representational structure harnessed by RL (***Figure 1***).

**Figure 1:**
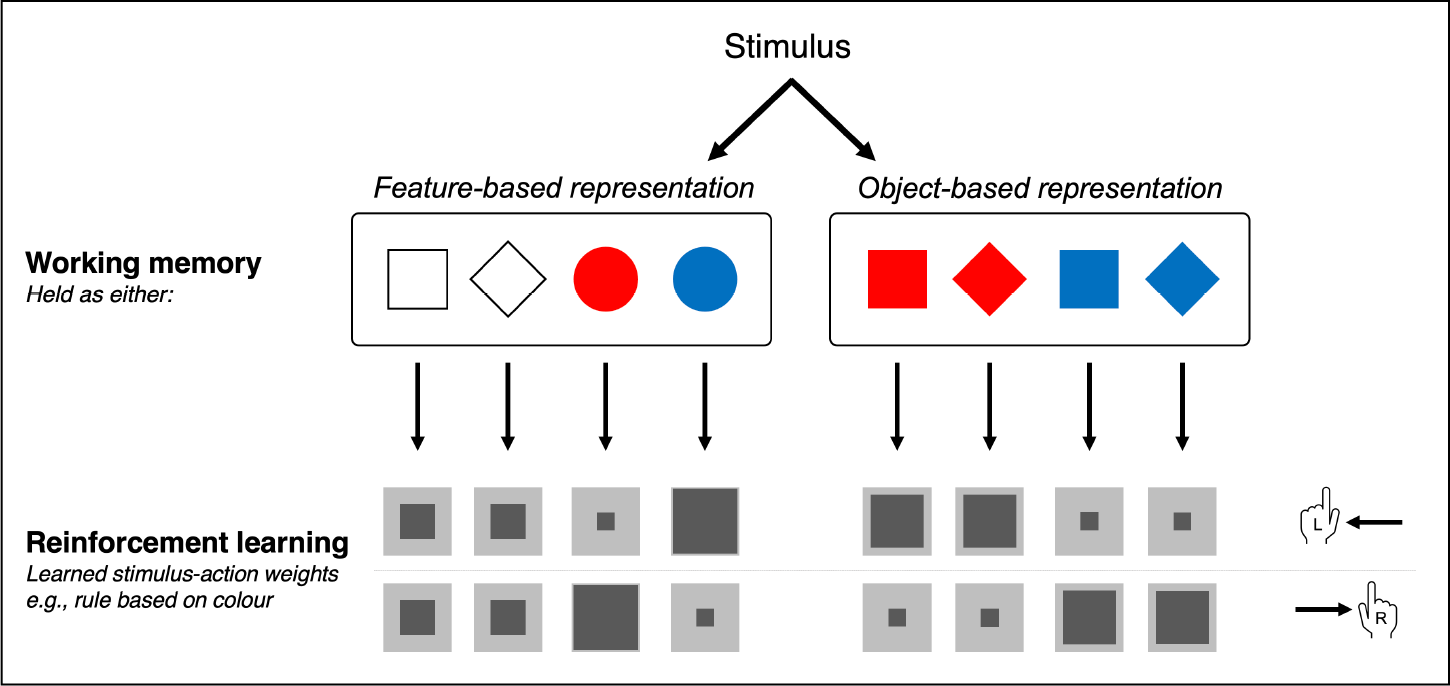
Proposed framework by which RL harnesses representations defined in WM. WM stores representation templates as feature-based or object-based representations. These representations are used in RL. The size of the dark grey box indicates the size of the learned stimulus-action weights, sometimes termed ‘Q-values’. Here, learnt weights for where colour define the rule is represented, where blue is associated to left and red is associated to right. For example, the blue states with left actions are shown to have higher weighting compared to blue states for right actions.

WM is a system that provides a temporary buffer to manipulate small amounts of information at a time^3^. Previous work suggests that it can be used as a short-term store for stimulus-action pairs^4,5^. Consequently, both WM and RL may be independently deployed to guide rewarded behaviour. Previous studies have shown that WM is more useful for small numbers of associations, whereas RL is deployed for larger sets of associations^4^, pointing to a task-dependent trade-off between WM and RL^6^. Alternatively, an account for WM contributing to augmented expectations of reward in the RL system has been reported^7^. However, we propose that WM also contributes to RL by holding structured information about currently relevant distinctions in the environment, which might provide the *template* that determines which states contextualise RL. Crucially, WM encodes information in a flexible and goal-directed form, which potentially lends great power to RL’s requirement to assign values to relevant features of the stimuli. This process relies on selecting the right representational format. Here, we assessed whether this selection is influenced by the representational format of the stimulus information held in WM. This would indicate that RL and WM collaborate, rather than compete, in assigning value to actions in different states.

An important aspect of the representational format in WM is whether visual features of an object are bound together to form an integrated representation^8–10^, or are maintained separately as flat sets and bound indirectly by a shared location^11–15^. WM is a system that prioritises task-relevant information, and representational states may be dynamic, holding both feature-based and object-based representations^16,17^. Furthermore, tasks can be designed to bias subjects towards maintaining either features or bound objects^18^. As representational formats in WM may either be beneficial or detrimental depending on the context, WM representations must be dynamically aligned to task demands to optimise goal-directed outcomes.

In this pre-registered study, we leveraged the possibility that WM representations can be flexibly shifted to investigate whether the representational format of WM could impact learning. We manipulated whether **features** (colour or shape) independently defined the stimulus-action rules or whether **objects** (colour-shape combinations) defined the stimulus-action rules. Simultaneously, we biased participants towards holding the RL stimulus in WM as a combination of features or a bound object. Half of the RL blocks required learning of rules based on features; the other half required learning of rules based on objects. Concurrently, half of the blocks probed the memory of the previously seen RL stimuli as single-feature, the other half as whole-object probes. This paradigm therefore aimed to causally bias representations towards either features or objects and assess its impact on learning.

Critically, the object and feature WM probes had exactly matched WM capacity requirements. If biasing the format of WM representations alters learning outcomes, this would suggest that the RL process uses representations held in WM to achieve learning. Specifically, we hypothesised that if representations in WM are used in RL, the representational structure that matched the RL rule (feature representation for feature-based rules, and object representation for object-based rules) would improve learning. Therefore, if representational biasing is successful, feature representations would facilitate feature-rule learning and lead to an optimised reinforcement strategy. Conversely, object representations should facilitate object-rule learning. Overall, we aimed to evaluate the extent to which learning-independent shifts in representations can alter learning behaviour and assess the interaction between WM and RL.

## Results

Participants completed 4 blocks of 128 trials each. Each trial consisted of Stage 1 - RL and Stage 2 – WM (***Figure 2***). In Stage 1, a binary choice (left or right) was made in response to a centrally presented coloured shape. In Stage 2, the memory of the stimulus encountered in Stage 1 was probed. Some blocks biased WM representations towards feature representations and others towards object representations. We evaluated how learning strategy changed with this WM bias (*see Methods*).

**Figure 2.**
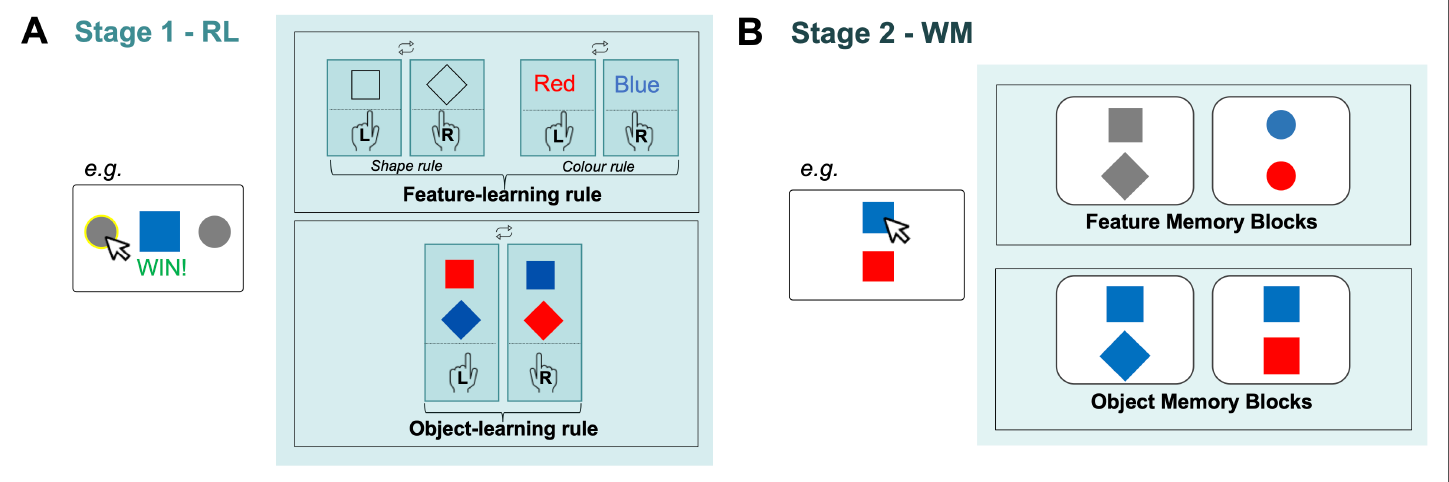
Experimental paradigm designed to assess impact of WM representational states on RL. **A) Stage 1:** Participants completed an RL task with stimuli involving colour and shape features. Half of the blocks had feature-based rules whilst the other half had object-based rules. In feature-based rule blocks, participants had to make action decisions (left or right) depending upon either shape or colour. In object-based blocks, the rules were based on conjunctions of features. The RL rule defining the stimulus action mappings changed every 32 trials. Reward was 80% probabilistic. **B) Stage 2:** After a 1.8s delay, participants recalled Stage 1 stimuli by selecting one of two items. In half of the blocks, they were probed with a single feature (just shape or just colour, top box), whilst in the other half, participants were probed using objects that contained both feature dimensions (bottom box). The memory demands were therefore equal in these two types of blocks. We quantified the impact of the Stage 2 WM representation on Stage 1 RL choice behaviour.

### RL and WM Accuracy

We first evaluated RL accuracy depending on rule type (feature or object rule) and WM probe type (feature or object probe). Both rule types were successfully learned over trials, (***Figure 3A***). There was higher average RL accuracy for feature rules than object rules (β = 0.33, *z* = - 8.43, *p* < .001, ***Figure 3B***) as well as better WM performance for feature memory probes than object memory probes (β = 0.20, *z* = 4.36, *p* < .001, ***Figure 3B***). The interaction between the RL rule and the WM probe was not significant (β = -0.10, *z* = -1.51, *p* = 0.13). Thus, contrary to our predictions, matching the memory probe method to the type of learning rule did not increase RL accuracy. These unexpected results challenge the initial hypothesis that matching the probe of the learning rule would enhance RL accuracy.

**Figure 3.**
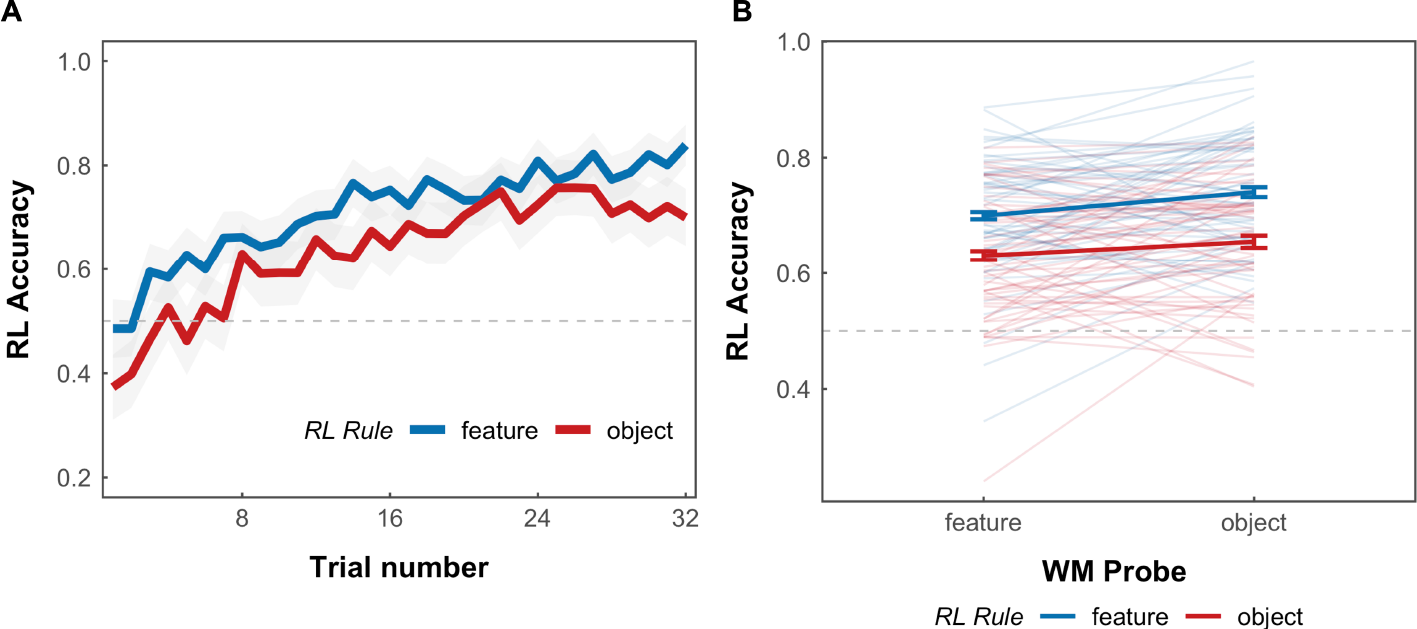
RL Accuracy Bas ed on RL Rule and WM Probe. A) Average performance for learning for each rule type for each trial. Each mini block consisted of 32 trials. Reward was 80% probabilistic, yielding an optimal performance of 0.8. The shaded area represents 95% confidence interval. B) Accuracy of RL stage split by RL rule, and by the type of WM probe in the WM stage revealing distinguishable patterns in RL accuracy based on rule and probe types. Individual line represents individual participants. Error bars are standard error of the mean. For both plots, the dotted line at 0.5 represents chance performance.

### Magnitude of reinforcement

To examine whether WM representations are used as a substrate of RL, we next investigated whether the effect of a reward on subsequent behaviour depends on the representational format defined in WM. We assessed the magnitude of reinforcement by quantifying p(stay), the probability of repeating an action (staying) when a reward was given, relative to when no reward was given (*see Methods*). This provided a metric to assess whether previous actions were reinforced by the previous reward or not. As expected, there was a strong effect of how many features changed from the previous trial on the likelihood of staying or switching choice based on the feedback provided in the previous trial (β = -1.07, *z* = -26.66, *p* < .001). We therefore split the trials depending on the numbers of feature changed. For trials where 2 features changed from previous, as expected, we found a significant effect of RL rule on p(stay) (β = 0.74, *z* = 9.50, *p* < .01) (***Figure 4*** right panel). We also observed that the WM probe affected p(stay) (β = -0.274, *z* = -2.90, *p* < .01). Crucially, a significant interaction effect between WM probe and RL rule was found (β = 0.36, *z* = 2.71, *p* < .01). Post-hoc comparisons revealed that when the WM probe was feature based, p(stay) was more optimal for feature rules (*z* = 2.91, *p* = -.02). In practice this meant that, when the rule was of the type “red = left, blue = right” or “square = left, diamond = right”, participants repeated their choice when the stimuli completely changed after reward versus no reward, but this learning was significantly stronger in blocks when the stimuli was probed in terms of individual features, compared to as a whole object. However, an object-based representation did not enhance p(stay) optimality for object rules (*z* = -0.99, *p* = - 0.76). The observed increase in p(stay) optimality for feature RL rules when paired with feature-based WM probes suggests that participants adapt reinforcement strategies more efficiently by aligning relevant feature representations in WM with task-specific RL rules.

**Figure 4.**
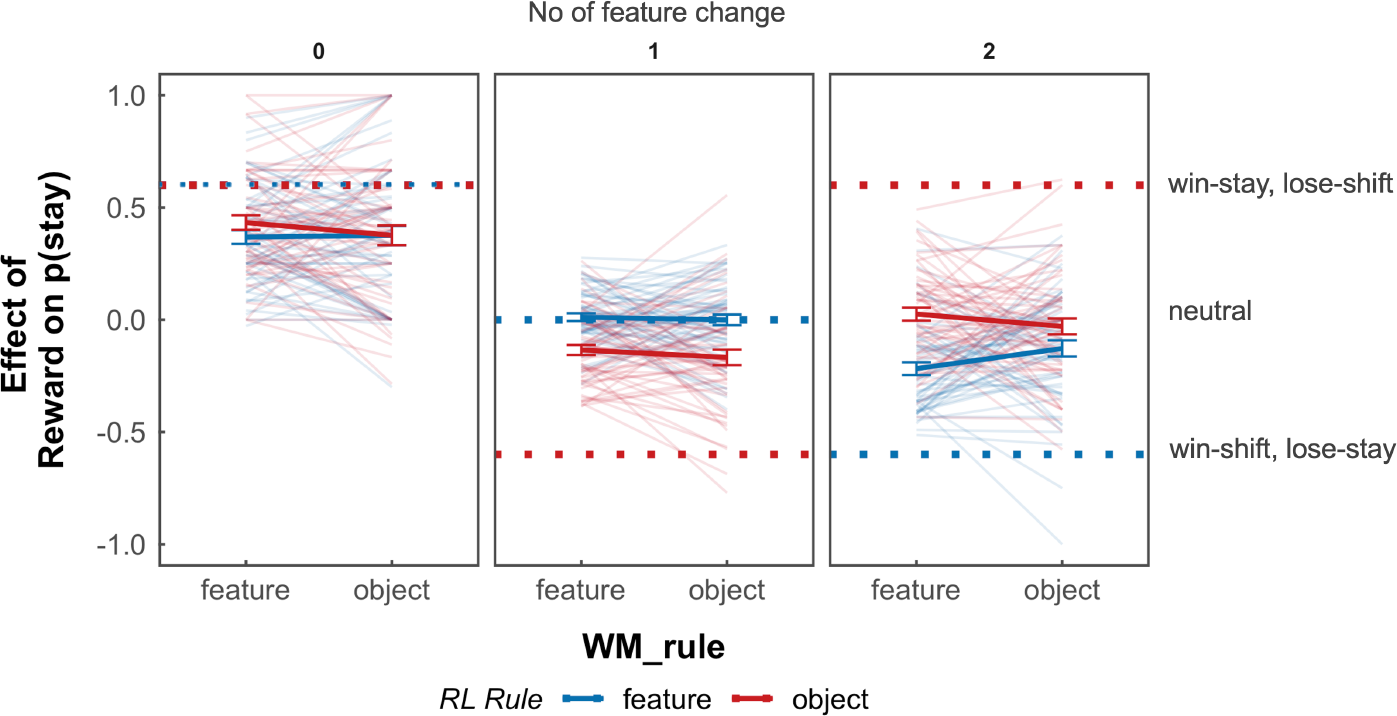
P(stay) by number of feature changes during RL trials. Left panel: When rewarded on RL trial t and no features change on RL trial t + 1, optimal behaviour is to maintain the previous choice, irrespective of the rule type. Middle panel: When rewarded on RL trial t, and one feature changes on RL trial t + 1, optimal p(stay) behaviour varies with rule types. For an object rule, the optimal behaviour is to switch, whereas for a feature rule, the decision depends on which feature changed. For instance, if the relevant feature changes, staying might be optimal, while if the irrelevant feature changes, switching might be more advantageous. Right panel: Optimal behaviour for two feature changes differs based on the RL rule. For object rules, individuals should stay, while for feature rules, individuals should switch. Optimality is represented by dotted lines. Individual lines represent individual participants. Error bars are standard errors of the mean. The text on the right describes the first-order strategy corresponding to the dotted lines.

### Exploratory analysis

We explored whether the effects of WM representational format were different at the start of the block after a rule change compared to later in the block. We found that in the first 16 trials, feature representations did not facilitate feature learning (*z* = 0.13, *p* = .72) nor did object representation facilitate object rule learning (*z* = -0.24, *p* = .29). The reported effects of feature representations facilitating feature learning were driven by learning strategies adopted in the last 16 trials (*z* = -0.42, *p* = .027). As per the previous main results, no effect was found for the object representations facilitating object rules, even in the last 16 trials (*z* = 0.057, *p* = .98).

We also assessed whether the type of feature being probed would affect p(stay). Specifically for feature rules where we could assess the congruency of rule and dimension probed, we assessed whether congruency improved p(stay) optimality. For example, if the rule was based on colour, and the WM probe on that given trial also tested memory for colour, would learning be stronger? No significant benefit of congruency was observed on p(stay) (β = 0.027, *z* = 0.048, *p* =.57) based on congruency when filtering out just the feature rules and dividing the analysis into congruent and incongruent trials. Further post-hoc tests did not show a benefit of congruent trials compared to incongruent trials (*t*(54) = 0.013, *p* = 0.99, *d* = 0.001). This showed that it was not simply the visual interaction of rule matching stimuli alone that optimised choice behaviour.

## Discussion

We proposed that working memory (WM) acts as a flexible store that represents aspects of visual input in a way that is relevant for reinforcement learning (RL). We expected that the format of WM will bias RL strategies across varying rule types (***Figure 1***). Our pre-registered experimental paradigm interleaving a WM task within an RL task showed that WM representations influence RL strategies. Crucially, instead of probing WM directly at the time of retrieval, we indirectly examined the *use* of WM contents in RL, which we argue interrogates the encoding and maintenance period and its use as a substrate for learning. This sheds light on how internal representations in WM might be harnessed in the RL process.

Whilst we expected that the WM probe type (feature or object) matched to the RL rule (feature or object rule) would enhance RL accuracy, our findings showed no benefit in overall performance accuracy. Accuracy is measured relative to a fictional ‘correct’ choice in a probabilistic task, that is inaccessible to the participant. Therefore, it is an impure measure of RL. We therefore assessed the trial-wise probability of repeating an action after receiving a reward. This metric, p(stay) varied with the type of WM probe. Participants adopted more optimal reinforcement strategies when relevant feature representations in WM were aligned with feature-specific RL rules. This is consistent with the hypothesis that when WM represented the stimuli as features, feature rules are easier to learn, compared to when WM represented the stimuli as objects. However, this is not the case for the opposite direction: object-based WM representations did *not* facilitate learning of object-specific RL rules.

One explanation for this asymmetry is that the preferred WM format for RL is features, which the feature WM probe accentuated. It can be speculated that object binding requires more effort, reducing the capacity for WM representations to influence RL. Alternatively, our WM bias may not have been strong enough. The cost of holding two features remains lower than the generally accepted limits of WM capacity and therefore the bias towards representations as a bound object may not have been successful. It is important in the future to move beyond lab-based studies using simple, static visual input with minimal ecological validity, and use tasks with richer and more complex environments with higher WM load^19^. Introducing greater complexity and uncertainty and reducing the precision of momentary internal representations could preferentially favour object representations. Finally, WM is not a unitary system^20^, and the same could also be said for RL. Thus, it is possible that the dovetailing of these two systems may not be complete, with only certain aspects of WM accessible for RL learning. For example, recent neural models of WM have distinguished sensory representations from “conjunctive” representations that enable this information to be bound into unified objects^21^.

Thus, one explanation for why only feature-based encoding influenced RL is because it is easier for the sensory component of WM representations to impact RL. In support of this explanation, an fMRI study found that post-reward activation of sensory regions was influenced by attention, whereby attending to faces increased BOLD signal in the fusiform face area after a reward^22^. This study also found that these BOLD signal changes in sensory regions were influenced by connectivity with reward-related regions such as the ventral striatum. Cumulatively, this suggests that sensory regions that support memory and RL may be inextricably linked, which is more strongly associated to single feature representations.

Our exploratory analysis revealed that probing one feature dimension of the stimulus, e.g. its colour, did not bias the learning towards that particular feature. Thus, simply attending to a feature did not facilitate RL, which speaks to the effect of representational formats in affecting RL. This may be because this attentional selection occurs *after* the reinforcement takes place. In contrast, the representational format (i.e. object vs features) remained constant for a whole block and could therefore bias both encoding and reinforcement. It remains an open question as to whether the representational bias is related only to how the stimulus is *encoded* into WM, or whether it can be subsequently affected by *changes* in WM representation, e.g. at the time of reinforcement.

Our present study cannot be explained simply in terms of attentional set, as only the *format* of the information changed, which was unrelated to the learning. Attentional set has traditionally been studied to assess how humans and animals select which dimensions of the stimulus are used (e.g. colour vs shape) for learning^23,24^. The Wisconsin card sorting task (WCST) requires individuals to sort cards based on changing rules without explicit instructions and has been used to study set-shifting and cognitive flexibility^2^. Rule learning requires correct credit assignment as well as maintenance and manipulation of information in WM. Similarly, in the Intra-dimensional/Extra-dimensional shift test (ID/ED)^25^, participants are required to shift attention between different dimensions of a stimulus. Extra dimensional shifts require attention to a previously irrelevant dimension, whereas intra dimensional shifts require attention to the same dimension. Like the WCST, the ID/ED task requires rule learning based on feedback and shifting the state on which you are informing your decisions is critical. Importantly, WM and set-shifting has been reported to act cooperatively in overlapping regions of the prefrontal cortex^26^. In this study the aspects of the stimulus that are relevant remain constant, so the effect we observed cannot be explained in terms of an effect on selecting or filtering the features, as has been done in these previously reported paradigms.

Finally, we have framed the effect of WM in terms of influencing RL, which suggests that value updates are applied only to the representations in WM. For example, if items are held as features, then RL updates the Q-values, which represents the expected future rewards for taking an action in a given state of actions, for feature states, i.e. States = {square, circle, red and blue}; whereas if items are held as objects, then RL updates the Q values of actions for object states, i.e. States = {red square, red circle, blue square, blue circle} (***Figure 1***). An alternative possibility is that in all cases, RL updates the values for some or all of these Q values, in a way that does not depend on WM. Then at the time of decision, the Q values are combined and weighted according to the contents of WM. In other words, WM may affect the value integration at the time of *decision-making*, rather than affecting learning directly. A similar dissociation has been proposed for model-based learning vs decision-making^27^. The current study cannot rule out this possibility and future studies are required to directly assess this.

Overall, our results underscore the interplay between WM representations and reinforcement learning strategies. We highlight the differential impact of WM representations on RL: “States” for reinforcement learning depend on the manner in which WM contents are represented. This may be the underlying mechanism for how biological reinforcement learning can be so flexible and context dependent. Hence, it could provide clues to enhancing learning in artificial agents. These findings prompt further exploration into the dynamic nature of WM in decision-making and the utilisation of internal representations defined in WM in other cognitive processes.

## Materials & Methods

### Participants

Data from 55 participants (mean age = 28.76, *SD* = 4.66 mean years of education = 15.53, *SD* = 3.40, 26 males: 29 females) was used for this study. The sample size was predetermined and pre-registered to achieve 0.90 power at a significance level of 0.05, to detect an effect size *d*’ of 0.40. This effect size was based on pilot data using a similar paradigm documented in the preregistration at https://osf.io/k7zjd (Shibata et al., 2023). Participants were recruited on a rolling basis until 55 individuals completed the following three criteria: 1) completed all experiment trials, 2) achieved an average asymptotic RL performance of 60% during the last 10 trials before a rule reversal (i.e. change in rules) and 3) responded to at least 75% of the WM trials correctly. A total of 84 people were initially recruited in the study and received monetary compensation. Thirty-one were excluded from the analysis as they did not meet the RL learning criterion. Three of those also had not attained the WM accuracy cut-off. No participants who performed above the RL performance threshold had a WM accuracy below 75%. All participants were fluent English speakers and reported no history of neurological or psychiatric illness. Informed consent was obtained from all participants through an online questionnaire (IRAS ID: 248379. Ethics Approval Reference: 18/SC/0448).

### Experimental tasks

#### Apparatus

The task was programmed using PsychoPy (V2022.2.4) and hosted on Pavlovia. Participants were recruited on the Prolific platform. Participants completed the task on their personal computer devices (participation through mobile and tablet devices was disabled) and were instructed to position themselves at arm’s length from their screen for the duration of the study.

#### Procedure

Participants completed a total of 512 trials, distributed across 4 blocks of 128 trials each. Each trial consisted of two distinct stages:

- *Stage 1 - RL* was a binary choice task which involved making a right or left click depending on the centrally presented coloured shape to receive a reward. Stimulus-response mappings, i.e. the rule defining the rewarded action corresponding to a particular stimulus, changed throughout the experiment.
- *Stage 2 – WM* was a probe for the memory of the stimulus they had just encountered in Stage 1. A blocked design allowed the biasing of WM representations into feature representations or object representations. We evaluated the impact of this bias on learning behaviour in Stage 1 - RL.

Prior to starting the main experiment, participants completed three practice blocks of 10 trials of each stage. The RL rule was provided for the practice blocks. Only individuals who achieved a performance accuracy of over 22/30 (73.33%) in both Stage 1 and Stage 2 trials during the practice were able proceed to the main experiment. This was implemented to verify task compliance and comprehension prior to starting the experiment. Participants were instructed to answer each trial as accurately and fast as possible. All participants were remunerated for their time.

### Stage 1 (*Figure 2A*)

The first part of the trial was a 2-choice RL task. Participants were presented with a stimulus appearing at the centre of the screen. This stimulus was made up of a combination of two features: colour (red or blue) and shape (square or diamond), making four possible stimuli: red square, red diamond, blue square, blue diamond. Participants were tasked with responding to a centrally presented stimulus with a mouse click on a grey circle either to the left, or to the right of the stimulus to win arbitrary rewards in the form points in a bank. A running total of points won was presented at the top of the screen throughout the task. No time limit was imposed. Visual feedback (‘Win!’ or ‘Lose!’) was instantly presented. A win resulted in an arbitrary reward of +5 points, whereas no points were awarded for losing.

The rewarded action was determined by an underlying RL rule that changed every 32 trials. As there were 128 trials per block, three rule reversals occurred within a single block. In half of the blocks, RL rules were based on individual features (*feature rules*), where either the colour or shape of the object was associated with reward. The other half of the blocks had rules based on objects (*object rules*), where the conjunction of colour and shape was associated with reward. The object-based rules never overlapped with feature-based rules such that object-based rules were always crossed. i.e. blue diamond, red square = left; red diamond, blue square = right OR red diamond, blue square = left; blue diamond, red square = right. Reward was 80% probabilistic, whereby for every 5 correct choices, only 4 were rewarded.

### Stage 2 (*Figure 2B*)

The second part of each trial consisted of a memory recall of the stimuli in Stage 1, after a 1.8s delay. Participants were asked to select which of the two stimuli, presented above and below the screen centre, corresponded to that observed in Stage 1. No time limit was imposed. In half of the blocks, participants were probed on one of the two features (either by colourless shapes or shapeless colours). Whether the shape or colour was probed was interleaved within a single block so that participants had to encode both features into working memory. In the other half of the blocks, participants were probed on objects with two features: coloured shape. The lure object could be different from the stimulus in either one or both features. Deterministic feedback and an arbitrary reward of +1 was provided after every choice. No reward was given for an incorrect response.

### Quantifying RL

We investigated reinforcement strategies in Stage 1 by examining the first-order strategy: the tendency to repeat an action, as a function of what was seen on the previous trial and whether the participant was rewarded. These simple heuristics allow us to quantify the type of representation subjects use for RL. We assessed whether the previous action was reinforced by the previous reward, i.e. the tendency to apply a “win-stay” or “lose-switch” strategy. To quantify this staying or switching behaviour, we calculated p(Stay), the probability of repeating an action (staying) when a reward was given, relative to when no reward was given as the following equation:

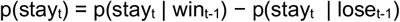

where t = trial number. A positive P(Stay) value indicates a higher likelihood of repeating an action, while a negative P(Stay) value suggests a greater probability of switching action.

Different choice strategies exist depending on the number of feature changes from one RL trial to the next. These strategies, illustrated in ***Figure 4***, inform the likelihood of repeating an action (p(stay)) under varying conditions.

- When successive trials are identical (no feature changes), there should be a strong positive reinforcement to repeat an action for both feature and object rules.
- If one feature from the previous trial is repeated, this leads to a neutral impact on action repetition for feature rules, but for object rules, it should prompt a switch in action.
- In cases where both features change from the previous trial, for feature rules the optimal action is to switch the action, but for object rules, the optimal action is to stay on the previously taken action.

We specifically assess choice behaviour on on trials with changes in both features as this trial type exhibits the most pronounced differences in p(stay) behaviour between the two rule types, which is a key point of interest in our analysis.

### Data pre-processing

All trials with a RT exceeding 3SD of the mean were considered outliers and removed from the analysis. Additionally, incorrect trials in the WM stage were removed as this indicated that the RL stimulus was not attended to (3.3% of otherwise retained trials). The steps of trial and participant exclusions are reported in the pre-registration. We additionally removed trials with both feature and shape probe dimensions, as these trials contain more information in the WM stage and do not act as a valid comparison.

### Statistics

Data was analysed using *R* (Rstudio 2022.12.0+353) and Matlab R2020b. A logistic mixed-effects regression model was fitted using the ‘glmer’ function from the ‘lme4’ package in *R*. The model aimed to evaluate the impact of RL Rule (feature or object) and WM Probe (feature or object) on binary RL choice outcomes (correct/incorrect). Specifically, the model investigated the individual contributions of RL Rule and WM Probe as well as their potential interaction effect on performance. Random effects were introduced for individual subjects to accommodate variability between conditions within each subject, accounting for within-subject correlations in the data. The model equation was formulated as follows: Accuracy (correct/incorrect) ∼ 1 + RL Rule * WM Probe + (1 | Subject). We also ran an additional mixed-effects model on p(stay) when two features changed from the previous trial: P(Stay) (stay/switch) ∼ 1 + RL Rule * WM Probe + (1 | Subject). We used the Tukey method for post-hoc comparisons. For between-subject exploratory analysis, Pearson correlations were used to test for correlation between variables. An alpha of .05 was used to report statistical significance and Greenhouse-Geisser correction was applied to degrees of freedom to correct for non-sphericity where appropriate.

## Code & Data Availability

The code and datasets analysed in the current study are available from the corresponding author on reasonable request.

## Author Contributions

S.G.M and K.S designed, interpreted and pre-registered the study. K.S acquired data and wrote the initial draft of the manuscript. V.K created the online version of the paradigm. K.S and V.K analysed the data with the help of S.G.M. S.G.M and S.J.F were involved in the initial conceptualisation of the idea. M.H and S.J.F provided critical feedback on the manuscript. All authors contributed to the final manuscript.

## Open Access Statement

For the purpose of Open Access, the author has applied a CC BY public copyright licence to any Author Accepted Manuscript (AAM) version arising from this submission.

